# Effect of D128N mutation on OsSERK2 in Xa21 mediated immune complex: an *in-silico* study

**DOI:** 10.1101/2022.11.30.518629

**Authors:** Raghib Ishraq Alvy, M H M Mubassir

## Abstract

Receptor-like kinases (RLKs) are plant proteins that form signaling circuits to transduce information through the plant cell membrane to the nucleus and activate processes that direct growth, development, stress response, and disease resistance. Upon sensing various environmental stress stimuli, RLKs interact with specific targets and recruits several other proteins to initiate the defense mechanism. Among many RLK subfamilies, leucine-rich repeat RLKs (LRR-RLKs) are the largest. Xa21, a member of LRR-RLK, is a vital receptor protein in rice plants that binds with bacterial RaxX21-sY, whereas OsSERK2 is a somatic embryogenic receptor kinase (SERK) acts as a coreceptor. This study focuses on the effect of a substitution mutation of aspartate128 with asparagine128 (D128N) in OsSERK2 on the interdependent binding pattern of the mentioned Xa21, RaxX21-sY, and OsSERK2 D128N proteins. The results showed that the D128N mutation in OsSERK2 can significantly change the interaction pattern of the critical residues of the OsSERK2 and affects its receptor-ligand (Xa21-RaxX21-sY) interaction in the complex.

## 1. INTRODUCTION

As multi-cellular sessile organisms that must respond to dynamic environments, plants need to effectuate and react toward numerous internal signals (Clark et al., 2001) to achieve their growth and metamorphosis (Prithiviraj et al., 2003; Sun et al., 2010), as well as to distinguish abundant input signals from their surroundings (Volkov & Ranatunga, 2006). Most of these signaling cues are detected on the cell periphery (Böhm et al., 2014; Kim et al., 2009), and plants have developed an idiosyncratic group of receptor-like kinases (RLKs) that can transmit extracellular signals throughout the membranes (De Smet et al., 2009; Shiu & Bleecker, 2001). RLKs are defined as a combination of a signal peptide, an extracellular domain, a transmembrane domain, and a cytoplasmic kinase domain together with the serine/threonine consensus sequence (Wang et al., 2007). In *Arabidopsis thaliana*, the number of leucine-rich repeat RLKs (LRR-RLKs) is greater than 600, originating from 13 subfamilies (Diévart & Clark, 2004; Shiu et al., 2004; Wang et al., 2013; Zhang, 1998; Zhang et al., 2006). These LRR-RLKs bind to somatic embryogenesis receptor kinases (SERKs) and form dimers (Mubassir; Schulze et al., 2010). The binding of SERKs with their respective LRR-RLKs is either ligand-dependent or independent. The best characterized SERK protein is BRI1-associated kinase 1 (BAK1), which binds with LRR-RLK FLS2, EFR, and BRI1. It forms a heterodimer, where ligand binding induces this heterodimerization (Chinchilla et al., 2007; Li et al., 2002; Schulze et al., 2010). BAK1 may also bind with RLK7 (Chowdhury & Mubassir, 2022; Shen et al., 2020), CORE (Jawad; Wang et al., 2016), and other similar receptors and thus become a common SERK coreceptor protein for many other LRR-RLKs and plays a key role as a regulator in plants’ innate immunity (Heese et al., 2007).

Among the LRR-RLKs, Xa21 is an important receptor protein of rice plants that binds with SERK OsSERK2 and forms a heterodimer. The Xa21 ectodomain contains 23 LRRs, a single transmembrane domain with a kinase domain followed by one juxtamembrane (JM) domain (Mubassir et al., 2020; Song et al., 1995). RaxX21, a bacterial peptide secreted by *Xanthomonas oryzae (Xoo)*, acts as a ligand for Xa21, the ectodomain of which binds with this pathogen-associated molecular pattern (PAMP) molecule. *Xoo* is the causative agent for bacterial leaf blight disease in the rice plant (NIÑO□LIU et al., 2006; Swings et al., 1990). The gene *XA21* confers resistance to the multiple isolates of *Xoo* and shows genetic and phenotypic diversity with the other rice plants (MHM Mubassir, Khondoker M Nasiruddin, Nazmul Hoque Shahin, Shamsun Nahar Begum, Manas Kanti Saha, & AQMB Rashid, 2016; MHM Mubassir, Khondoker M Nasiruddin, Nazmul Hoque Shahin, Shamsun Nahar Begum, Manas Kanti Saha, & AQM Bazlur Rashid, 2016; M Mubassir et al., 2016; Wang et al., 1996).

When RaxX21 is secreted, sulfation occurs in its tyrosine region (RaxX21-sY) (Pruitt et al., 2015), which, as a result, gives more stability to the bacterial peptide (Mubassir et al., 2017). After RaxX21-sY binds with the ectodomain of Xa21, it recruits co-receptor SERK protein OsSERK2 and activates the defense signal by forming a heterodimer structure (Chen et al., 2014; Mubassir et al.). Interestingly the Xa21 also can interact with its coreceptor OsSERK2 without the presence of RaxX21-sY (Chen et al., 2014). Besides associating with Xa21, OsSERK2 also binds with other rice proteins such as OsBRI1, which is important for brassinosteroid-regulated development, as well as OsFLS2; and Xa3, which are important for initiating the defense mechanism in rice plants (Chen et al., 2014; Park et al., 2011).

In rice plants, OsSERK2 acts as a functional homolog of BAK1, transphosphorylating the kinase domains of these rice receptor proteins (Chen et al., 2014; McAndrew et al., 2014). Additionally, the structure of OsSERK2 is highly similar to BAK1 coreceptor. The short-curved solenoid ectodomain of OsSERK2, composed of an N-terminal LRRNT and five LRRs, has a 67% similarity with the BAK1 (McAndrew et al., 2014; Santiago et al., 2013). Moreover, the binding pattern of OsSERK2 and BAK1 reveals that both of the SERK coreceptors bind with their respective pattern recognition receptor proteins (PRRs) at the concave side of their ectodomains (Koller & Bent, 2014; Santiago et al., 2013; Sun, Han, et al., 2013; Sun, Li, et al., 2013).

The mutation in BAK1 of Asp122 to asparagine alters its interaction with its respective PRRs (Jaillais et al., 2011). A recently produced crystallographic structure of FLS2-BAK1 and BRI1-BAK1 has revealed that, although mutation in Asp122 alters the overall interaction, there is no direct contact of this particular residue with FLS2 or BRI1 (Santiago et al., 2013; Sun, Han, et al., 2013). Moreover, this residue was not predicted to be glycosylated in the D122N mutant of BAK1 (Chauhan et al., 2012; McAndrew et al., 2014), which suggests that mutation in this particular residue indirectly influences binding by altering the position of residues near it.

Asp128 residue of OsSERK2 forms hydrogen bonds with Ser126 and a salt bridge with Arg152. Mutation in Asp128 alters the binding of this residue with Arg152; in this case, Arg152 interacts with the residue Glu174 (McAndrew et al., 2014). This aspartate is conserved among all the rice SERK proteins and *A. thaliana*. Asp128 in rice OsSERK2 is the corresponding residue of BAK1 and is located in the LRR3 region of OsSERK2 (McAndrew et al., 2014). However, the impact of this mutation in the Xa21 LRR-RaxX21-sY-OsSERK2 LRR complex is yet to be elucidated. In this study, we employed a detailed in silico approach to analyze the impact of the Asp128 mutation in OsSERK2 in the Xa21-mediated immune complex. We showed how this mutation affects the interaction pattern of the residues of OsSERK2.

## 2. RESULTS AND DISCUSSION

### 2.1 Changes in the behavior of the mutated residue

We first checked our designed OsSERK2 mutated structure with the crystal OsSERK2 D128N (4q3i) structure and found both structures similar (Fig. S1). We then proceed with our designed structure to do the molecular dynamics simulation. The previously solved crystal structure of OsSERK2 showed that due to this mutation, the salt bridge interaction of Arg152 with Asp128 was demolished (McAndrew et al., 2014). Our study also found the same phenomena (Fig. 1a-1b). We observed that, Arg152 of OsSERK2 D128N shifted its salt bridge interaction with Glu174 (Fig. 1c) (Table S1), which also supports the previous structural study (McAndrew et al., 2014). Again, we noticed Ser126 had a hydrogen bond with Asp128 in OsSERK2 (Fig. 1b), but in the case of Os-SERK2 D128N, that interaction was discontinued (Fig. 1c), which also justifies the previous study (McAndrew et al., 2014). These changes in intra-protein interaction might lead to several changes in the interaction of the prominent residues of the complex of OsSERK2 with its PRR Xa21 and respective PAMP RaxX21. Previous studies have shown that a similar mutation in BAK1 (*BAK1 elg*, which results in aspartate 122 substitutions with asparagine) showed no difference in its interaction with its receptor BRI1 (Jaillais et al., 2011). However, in a recent structural and biochemical study, it was observed that this mutation can disrupt BAK1’s ability to interact with the ectodomain of BRI1 receptor pseudo kinases (Hohmann et al., 2018) and can lead to the stabilization of certain SERK3 residues (Hohmann et al., 2018; Santiago et al., 2013). Moreover, this mutation could also lead to the impairment of its (*BAK1 elg*) ligand-induced association with FLS2 (Jaillais et al., 2011).

**Fig 1.**
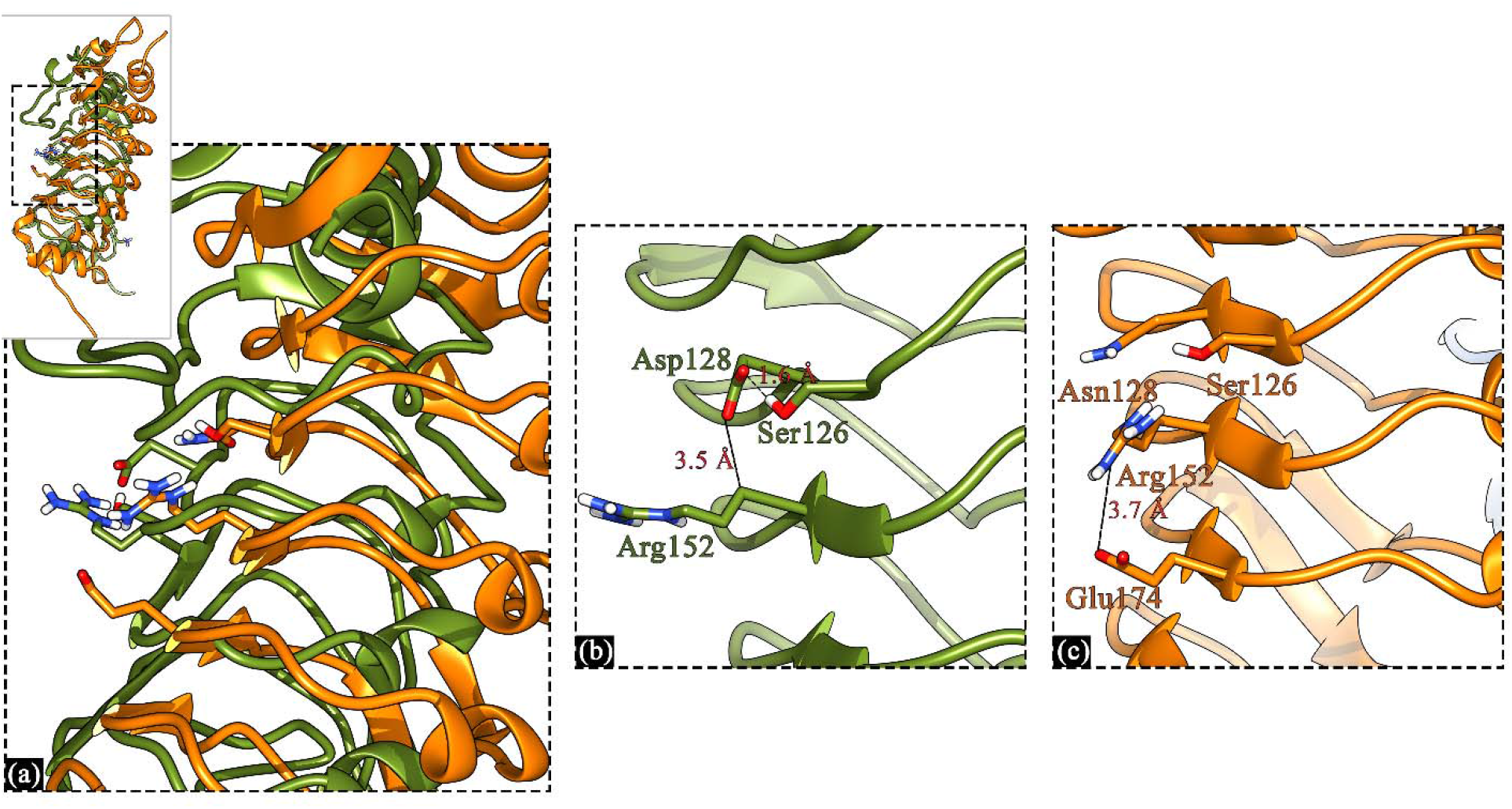
Changes in the interaction of the mutated residue with the neighboring residues. (a) The superimposed cartoon structure of OsSERK2 and OsSERK2 D128N focused on the mutated area. (b) Cartoon structure of OsSERK2; where Asp128 formed an s hydrogen bond with Ser126, and a salt bridge interaction with Arg152 of itself. (c) Cartoon structure of OsSERK2 D128N; where mutated residue Asn128 has no interaction with Ser126 and Arg152. Rather Arg152 formed a salt bridge interaction with Glu174 (Table S1). Cartoon: OsSERK2 LRR (green), OsSERK2 D128N LRR (orange); Stick: Asp128/Asn128, Arg152, Ser126, Glu174; Both structures were evaluated after 100ns MD simulation.

### 2.2 Changes in interactions of prominent residues of the proteins

Our previous study showed that Arg185 and Arg230 from Xa21, Val2, and Lys15 from RaxX21-sY, and Lys164 from OsSERK2 act as prominent residues for the wild Xa21 LRR-RaxX21-sY-OsSERK2 LRR complex (having wild OsSERK2 LRR) (Mubassir et al., 2020). For this wild complex, Arg185 of Xa21 makes a hydrogen bond with Leu52 of OsSERK2, and Arg230 makes a hydrogen bond with Asp6 of RaxX21-sY within 3.5 A (Fig. 2a). Moreover, Val2 and Lys15 of RaxX21-sY form hydrogen bonds with Asn331, and Cys382 of Xa21, respectively (Fig. 2b). Lys164 of OsSERK2 interacts with Asp565 of Xa21 by forming an ionic bond within 4Å (Fig. 2c) (Mubassir et al., 2020).

**Fig 2.**
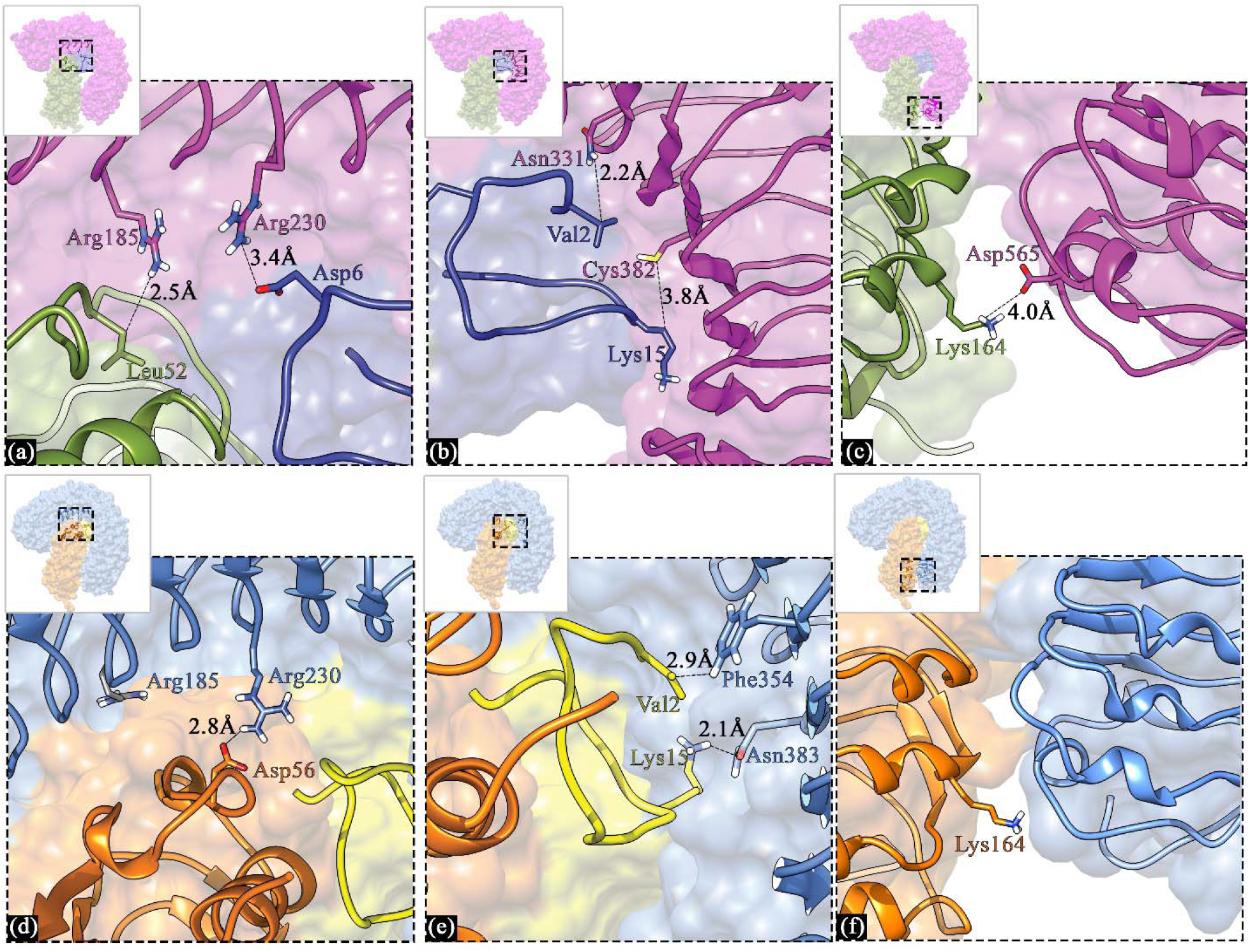
Changes in interactions of prominent residues of the complex. (a) Arg185 and Arg 230 of Xa21 form hydrogen bonds with Leu52 of OsSERK2 and Asp6 of RaxX21-sY in the wild complex. (b) Val2 and Lys15 of RaxX21-sY form hydrogen bonds with Asn331 and Cys382 of Xa21 in the wild complex. (c) Lys164 of OsSERK2 forms an ionic bond with Asp565 of Xa21 in the wild complex. (d) Arg185 of Xa21 discontinues the previous interaction, and Arg230 of Xa21 forms a new hydrogen bond with Asp56 of OsSERK2 D128N in the mutated complex (Table S2). (e) Val2 and Lys15 of RaxX21-sY form a new hydrophobic interaction with Phe354 and hydrogen bond Asn383 of Xa21 in the mutated complex (Table S2 and Table S3). (f) Lys164 of OsSERK2 D128N shows no interaction within 4.0Å in the mutated complex. Cartoon: Xa21-RaxX21-sY-OsSERK2 complex where Xa21 (magenta), RaxX21-sY (blue), and OsSERK2 (green), and Xa21-RaxX21-sY-OsSERK2 D128N complex where Xa21 (light blue), RaxX21-sY (yellow), and OsSERK2 D128N (orange); Stick: prominent and interacting residues of both complex; all the complexes were investigated after 100ns MD simulation.

On the contrary, due to the D128N mutation in OsSERK2, these prominent residues changed their interaction patterns. The previous interaction of Arg185 and Arg230 of Xa21 got discontinued; instead, Arg230 of Xa21 established a new hydrogen bond with Asp56 of OsSERK2 D128N (Fig. 2d) (Table S2). Val2 from RaxX21-sY formed a hydrophobic interaction with Phe354 of Xa21, and Lys15 bonded with Asn383 of Xa21 by forming a hydrogen bond (Fig. 2e) (Table S2 and Table S3). No bond formation of Lys164 from OsSERK2 D128N is found within 4Å in this mutated complex (Fig. 2f). These changes in interactions of the prominent residues point to the overall interaction pattern in the complex due to the mutation. Our findings support the previous finding of the mutation effect on the similar protein BAK1, where it was observed that mutation in BAK1 hampers the interaction between BAK1 and BRI1 receptor pseudo kinases (Hohmann et al., 2018). Also, a mutation in BAK1 might lead to the impairment of its (*BAK1 elg*) ligand-induced association with FLS2 (Jaillais et al., 2011).

### 2.3 RMSD and RMSF of the Xa21 LRR-RaxX21-sY-OsSERK2 D128N LRR complex

The stability of the complex was measured in terms of deviations by analyzing the root mean square deviation (RMSD) after performing a 100ns molecular dynamics simulation. The backbone of the complex showed the least variable RMSD in the simulated system. It deviated from 0.0004 to 0.75 nm during the entire simulation period (Fig. 3a). The complex’s standard deviation (SD) was 0.05 nm. The average deviation of the complex is 0.49 nm, which indicates that the Xa21 LRR-RaxX21-sY-OsSERK2 D128N LRR complex is stable and favorable.

**Fig 3.**
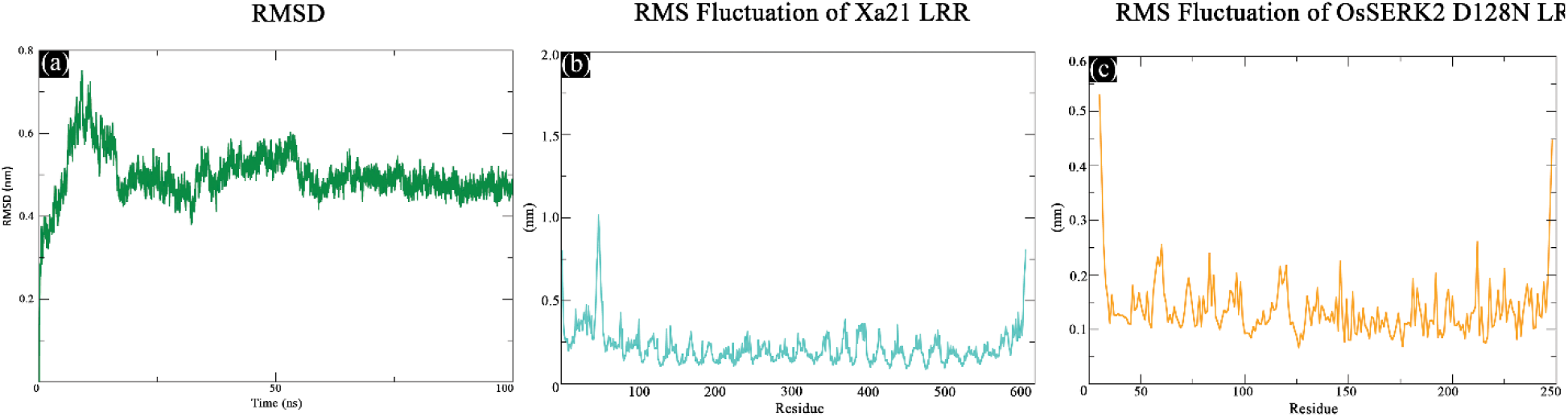
RMS deviation (RMSD) and RMS fluctuation (RMSF). (a) RMSD of the complex Xa21 LRR–RaxX21-sY–OsSERK2 D128N LRR from the 100ns trajectory. (b) RMS fluctuations of the residues of Xa21 LRR from the 100ns trajectory. (c) RMS fluctuations of the residues of OsSERK2 D128N LRR from the 100ns trajectory.

Moreover, we utilized Rosetta (Das & Baker, 2008) to relax and measure the energy of the after-simulated complexes, and it also aligns with the results we got. It showed a lower total score (1685.627 kJ/mol) for the mutated complex where fa_atr (Lennard-Jones attractive between atoms in different residues) contributed the most (−4728.319 kJ/mol) to minimize the energy (Table S4).

From the 100 ns MD trajectories, the RMSFs of the residues of Xa21 LRR, RaxX21-sY, and OsSERK2 D128N LRR were calculated. Analyses of the overall results revealed that most residues fluctuated by less than 0.21 nm for Xa21 LRR and 0.13 nm for OsSERK2 D128N LRR (Fig. 3b-3c). Moreover, the residues stated as prominent with low RMSFs, in our previous study [19], showed slightly higher in the mutated complex. For Xa21, Arg185 and Arg230, which showed a different binding pattern, exhibited high RMSF than the average RMSF of Xa21 LRR (Fig. 3b). Also, for RaxX21-sY, Val2 and Pro14, which were considered as important residues for binding with Xa21, showed similar phenomena. Furthermore, Lys164 of OsSERK2 D128N had a low RMSF with a value of 0.14 nm (Fig. 3c). On the other hand, the mutated residue Asn128 of OsSERK2 D128N showed a very low RMSF value (0.08 nm) alongside Arg152 and Ser126. The overall RMSD and RMS fluctuation of Xa21 LRR, RaxX21-sY, and OsSERK2 D128N LRR seem less fluctuating and more stable than the wild complex. These results also support the previous work where it was observed that mutation in the coreceptor could lead to the stabilization of certain SERK3 residues (Hohmann et al., 2018; Santiago et al., 2013)

## 3. MATERIALS AND METHODS

### 3.1 Mutating co-receptor OsSERK2

In our previous study, OsSERK2 LRR was docked with the PRR Xa21 LRR and PAMP RaxX21-sY (Mubassir et al., 2020). Multiple *in silico* modeling approaches were used to predict the 3D model of a full plant PRR protein, Xa21, and only the LRR part of Xa21 was examined for the interaction (Mubassir et al., 2020). OsSERK2 LRR (PDB ID: 4q3g) was obtained from Protein Data Bank (Burley et al., 2017; McAndrew et al., 2014). In the current study, OsSERK2 LRR of that complex was point mutated using rotamer tools of UCSF Chimera (Pettersen et al., 2004), where, Dunbrack library was used (Dunbrack Jr, 2002). The point mutation was performed to mutate 128th positioned aspartate into asparagine. The most probable form of asparagine (probability = 0.473) was selected for this position, and then a new PDB file was created for molecular dynamics (MD) simulation.

### 3.2 MD simulation of Xa21 LRR, RaxX21-sY, and OsSERK2 D128N LRR complex

The GROMACS software suite (version 5.1) (Van Der Spoel et al., 2005) was used to carry out the molecular dynamics (MD) simulation process of the complex Xa21 LRR-RaxX21-sY-OsSERK2 D128N LRR. GROMOS 54a7 (Schmid et al., 2011) united force field was applied for the simulation process. A cubic box with a distance of 1Å between the surfaces and edges of the complex was set for the solvation. The complex was then solvated using the SPC water model (Fuhrmans et al., 2010) using the *gmx solvate* tool, and *gmx genion* was used to neutralize the system, and then energy minimization was performed using the *gmx grompp* tool. Then the system was equilibrated for two ns NVT and one ns NPT, respectively, setting the temperature at 300 K and pressure at 1 atm. Finally, a 100 ns MD simulation was run for the system using the *gmx grompp* tool. To analyze the stability of the complex over the simulation period, the GROMACS *gmx rms* tool was used, and the GROMACS *gmx rmsf* tool was used to analyze individual residual fluctuations.

### 3.3 Investigation of the interaction of Xa21 LRR and RaxX21-sY with OsSERK2 D128N LRR

The binding pattern of OsSERK2 D128N LRR with its PRR Xa21 LRR and respective PAMP RaxX21-sY was examined using UCSF Chimera. To analyze different types of interactions alongside salt-bridge, Protein Interaction Calculator (PIC) (Tina et al., 2007) and Evaluating the Salt Bridges in Proteins (ESBRI) (Costantini et al., 2008) web tools were used. The Protein Interactions Calculator (PIC) website uses the coordinates of a protein’s or assembly’s 3D structure to calculate numerous interactions, including disulfide bonds, contacts between hydrophobic residues, ionic interactions, hydrogen bonds, aromatic-aromatic interactions, aromatic-sulfur interactions, and cation-interactions inside a protein or between proteins in a complex. Interactions are computed using standard and publicly available criteria. The convenience of accessing inter-residue interaction calculations in a single location is a benefit of employing a PIC server. It also calculates the available surface area and the distance between a residue and the protein’s surface. In contrast, ESBRI is a web-based software tool used for calculating salt bridge interactions of protein complex (Costantini et al., 2008).

### 3.4 Analysis of the mutated residue of OsSERK2 D128N LRR

Using the UCSF Chimera *clash and contact* tool, the interactions of Asp128 of Os-SERK2 and Asn128 of OsSERK2 D128N with their neighboring residues were observed, where atoms of the selected residues were designated. Also, by using the UCSF Chimera *distance measurement* tool, the distances between interactive atoms were visualized.

## 4. CONCLUSIONS

In this study, we tried to extensively investigate the effect of a D128N mutation in OsSERK2 in the Xa21-mediated immune complex. One of the key findings was the behavioral changes in the amino acid residues of the Xa21, RaxX21-sY, and OsSERK2 proteins due to the mutation. Arg185 and Arg230 of Xa21, which were identified as prominent residue in the previous study, change their interaction pattern in mutated complex, Val2 and Lys15 of RaxX21-sY also show similar phenomena, and Lys164 of OsSERK2 D128N discontinue previous interaction in the mutated complex, which significantly differs from the complex of Xa21 containing the wild type OsSERK2. In the wild type, Asp128 of OsSERK2 had two notable interactions, one was a salt bridge interaction with Arg152, and another was a hydrogen bond with Ser126. However, due to the D128N mutation of OsSERK2, Arg152 of OsSERK2 D128N established another salt bridge interaction with Glu174, and Ser126 discontinued the hydrogen bond.

Though with this *in-silico* study, we successfully showed the effect of D128N mutation in OsSERK2 itself and in the Xa21-mediated defense complex, a wet-lab structure-based approach (X-ray crystallography or cryogenic electron microscopy) is crucial to verify the data further. Also, an investigation of the changes in the phenotypic expression of the rice plant due to this mutation is pivotal. We firmly believe these findings will significantly contribute to those structures and phenotypic study and, therefore, can aid the scientific community in studying further the structural basis of Xa21-mediated immunity and, in general, the first layer of the plant defense mechanism.

## AUTHOR CONTRIBUTIONS

M.M. conceptualized and designed the study. R.I.A. performed the computational work and analysis. M.M. supervised the project. R.I.A. and M.M. co-wrote the first draft. All authors have read and agreed to the published version of the manuscript.

## ACKNOWLEDGMENTS

The authors thank Marzia Khatun, Tasfia Tawhid Supti, Ramen Chowdhury, and Tanvir Jawad for their continuous support throughout the research.

## REFERENCES

Böhm, H., Albert, I., Fan, L., Reinhard, A., & Nürnberger, T. (2014). Immune receptor complexes at the plant cell surface. Current opinion in plant biology, 20, 47–54.

Burley, S. K., Berman, H. M., Kleywegt, G. J., Markley, J. L., Nakamura, H., & Velankar, S. (2017). Protein Data Bank (PDB): the single global macromolecular structure archive. Protein Crystallography, 627–641.

Chauhan, J. S., Bhat, A. H., Raghava, G. P., & Rao, A. (2012). GlycoPP: a webserver for prediction of N-and O-glycosites in prokaryotic protein sequences. PloS one, 7(7), e40155.

Chen, X., Zuo, S., Schwessinger, B., Chern, M., Canlas, P. E., Ruan, D., Zhou, X., Wang, J., Daudi, A., & Petzold, C. J. (2014). An XA21-associated kinase (OsSERK2) regulates immunity mediated by the XA21 and XA3 immune receptors. Molecular plant, 7(5), 874–892.

Chinchilla, D., Zipfel, C., Robatzek, S., Kemmerling, B., Nürnberger, T., Jones, J. D., Felix, G., & Boller, T. (2007). A flagellin-induced complex of the receptor FLS2 and BAK1 initiates plant defence. Nature, 448(7152), 497–500.

Chowdhury, R., & Mubassir, M. (2022). How Arabidopsis Receptor-Like Kinase 7 (RLK7) Manifests: Delineating Its Structure and Function. Advances in Agriculture, 2022.

Clark, G. B., Thompson Jr, G., & Roux, S. J. (2001). Signal transduction mechanisms in plants: an overview. Current Science, 170–177.

Costantini, S., Colonna, G., & Facchiano, A. M. (2008). ESBRI: a web server for evaluating salt bridges in proteins. Bioinformation, 3(3), 137.

Das, R., & Baker, D. (2008). Macromolecular modeling with rosetta. Annu. Rev. Biochem., 77, 363–382.

De Smet, I., Voss, U., Jürgens, G., & Beeckman, T. (2009). Receptor-like kinases shape the plant. Nature cell biology, 11(10), 1166–1173.

Diévart, A., & Clark, S. E. (2004). LRR-containing receptors regulating plant development and defense.

Dunbrack Jr, R. L. (2002). Rotamer libraries in the 21st century. Current opinion in structural biology, 12(4), 431–440.

Fuhrmans, M., Sanders, B. P., Marrink, S.-J., & de Vries, A. H. (2010). Effects of bundling on the properties of the SPC water model. Theoretical Chemistry Accounts, 125(3), 335–344.

Heese, A., Hann, D. R., Gimenez-Ibanez, S., Jones, A. M., He, K., Li, J., Schroeder, J. I., Peck, S. C., & Rathjen, J. P. (2007). The receptor-like kinase SERK3/BAK1 is a central regulator of innate immunity in plants. Proceedings of the National Academy of Sciences, 104(29), 12217–12222.

Hohmann, U., Nicolet, J., Moretti, A., Hothorn, L. A., & Hothorn, M. (2018). The SERK3 elongated allele defines a role for BIR ectodomains in brassinosteroid signalling. Nature Plants, 4(6), 345–351.

Jaillais, Y., Belkhadir, Y., Balsemão-Pires, E., Dangl, J. L., & Chory, J. (2011). Extracellular leucine-rich repeats as a platform for receptor/coreceptor complex formation. Proceedings of the National Academy of Sciences, 108(20), 8503–8507.

Jawad, T. Bioinformatics Approach of Structural Modelling and Molecular Dynamics Simulation of Pattern Recognition Receptor CORE.

Kim, T.-W., Guan, S., Sun, Y., Deng, Z., Tang, W., Shang, J.-X., Sun, Y., Burlingame, A. L., & Wang, Z.-Y. (2009). Brassinosteroid signal transduction from cell-surface receptor kinases to nuclear transcription factors. Nature cell biology, 11(10), 1254–1260.

Koller, T., & Bent, A. F. (2014). FLS2-BAK1 extracellular domain interaction sites required for defense signaling activation. PLoS One, 9(10), e111185.

Li, J., Wen, J., Lease, K. A., Doke, J. T., Tax, F. E., & Walker, J. C. (2002). BAK1, an Arabidopsis LRR receptor-like protein kinase, interacts with BRI1 and modulates brassinosteroid signaling. Cell, 110(2), 213–222.

McAndrew, R., Pruitt, R. N., Kamita, S. G., Pereira, J. H., Majumdar, D., Hammock, B. D., Adams, P. D., & Ronald, P. C. (2014). Structure of the OsSERK2 leucine-rich repeat extracellular domain. Acta Crystallographica Section D: Biological Crystallography, 70(11), 3080–3086.

Mubassir, M. A Synopsis of Different Plant LRR-RLKs Structures and Functionality.

Mubassir, M., Naser, M. A., Abdul-Wahab, M. F., & Hamdan, S. A Brief Overview on Early Events of Xa21 Mediated Pattern Triggered Immunity.

Mubassir, M., Naser, M. A., Abdul-Wahab, M. F., & Hamdan, S. (2017). In-Silico Structural Modeling and Molecular Dynamics Simulation of Pathogen-Associated Molecular Pattern RAXX21. Journal of Chemical and Pharmaceutical Sciences, 10(1), 121–126.

Mubassir, M., Naser, M. A., Abdul-Wahab, M. F., Jawad, T., Alvy, R. I., & Hamdan, S. (2020). Comprehensive in silico modeling of the rice plant PRR Xa21 and its interaction with RaxX21-sY and OsSERK2. RSC advances, 10(27), 15800–15814.

Mubassir, M., Nasiruddin, K. M., Shahin, N. H., Begum, S. N., Saha, M. K., & Rashid, A. (2016). Morpho-molecular screening for bacterial leaf blight resistance in some rice lines and varieties. Journal of Plant Sciences, 4(6), 146–152.

Mubassir, M., Nasiruddin, K. M., Shahin, N. H., Begum, S. N., Saha, M. K., & Rashid, A. B. (2016). SSR Marker Based Genetic Diversity Analysis of Some Rice Lines and Varieties for Bacterial Leaf Blight Resistance. Journal of Pharmaceutical Chemical and Biological Sciences, 4(4), 475–486.

Mubassir, M., Nasiruddin, K. M., Shahin, N. H., Begum, S. N., Sultana, A., & Rashid, A. (2016). Measurement of Phenotypic Variation for Control and Bacterial Leaf Blight Inoculated Rice Lines and Varieties. Am. J. Biosci. Bioeng, 4, 59–64.

NiñO□Liu, D. O., Ronald, P. C., & Bogdanove, A. J. (2006). Xanthomonas oryzae pathovars: model pathogens of a model crop. Molecular plant pathology, 7(5), 303–324.

Park, H. S., Ryu, H. Y., Kim, B. H., Kim, S. Y., Yoon, I. S., & Nam, K. H. (2011). A subset of OsSERK genes, including OsBAK1, affects normal growth and leaf development of rice. Molecules and cells, 32(6), 561–569.

Pettersen, E. F., Goddard, T. D., Huang, C. C., Couch, G. S., Greenblatt, D. M., Meng, E. C., & Ferrin, T. E. (2004). UCSF Chimera—a visualization system for exploratory research and analysis. Journal of computational chemistry, 25(13), 1605–1612.

Prithiviraj, B., Zhou, X., Souleimanov, A., Kahn, W., & Smith, D. (2003). A host-specific bacteria-to-plant signal molecule (Nod factor) enhances germination and early growth of diverse crop plants. Planta, 216(3), 437–445.

Pruitt, R. N., Schwessinger, B., Joe, A., Thomas, N., Liu, F., Albert, M., Robinson, M. R., Chan, L. J. G., Luu, D. D., & Chen, H. (2015). The rice immune receptor XA21 recognizes a tyrosine-sulfated protein from a Gram-negative bacterium. Science advances, 1(6), e1500245.

Santiago, J., Henzler, C., & Hothorn, M. (2013). Molecular mechanism for plant steroid receptor activation by somatic embryogenesis co-receptor kinases. Science, 341(6148), 889–892.

Schmid, N., Eichenberger, A. P., Choutko, A., Riniker, S., Winger, M., Mark, A. E., & van Gunsteren, W. F. (2011). Definition and testing of the GROMOS force-field versions 54A7 and 54B7. European biophysics journal, 40(7), 843–856.

Schulze, B., Mentzel, T., Jehle, A. K., Mueller, K., Beeler, S., Boller, T., Felix, G., & Chinchilla, D. (2010). Rapid heteromerization and phosphorylation of ligand-activated plant transmembrane receptors and their associated kinase BAK1. Journal of Biological Chemistry, 285(13), 9444–9451.

Shen, J., Diao, W., Zhang, L., Acharya, B. R., Wang, M., Zhao, X., Chen, D., & Zhang, W. (2020). Secreted peptide PIP1 induces stomatal closure by activation of guard cell anion channels in Arabidopsis. Frontiers in Plant Science, 11, 1029.

Shiu, S.-H., & Bleecker, A. B. (2001). Plant receptor-like kinase gene family: diversity, function, and signaling. Science’s STKE, 2001(113), re22–re22.

Shiu, S.-H., Karlowski, W. M., Pan, R., Tzeng, Y.-H., Mayer, K. F., & Li, W.-H. (2004). Comparative analysis of the receptor-like kinase family in Arabidopsis and rice. The plant cell, 16(5), 1220–1234.

Song, W.-Y., Wang, G.-L., Chen, L.-L., Kim, H.-S., Pi, L.-Y., Holsten, T., Gardner, J., Wang, B., Zhai, W.-X., & Zhu, L.-H. (1995). A receptor kinase-like protein encoded by the rice disease resistance gene, Xa21. science, 270(5243), 1804–1806.

Sun, Y., Fan, X.-Y., Cao, D.-M., Tang, W., He, K., Zhu, J.-Y., He, J.-X., Bai, M.-Y., Zhu, S., & Oh, E. (2010). Integration of brassinosteroid signal transduction with the transcription network for plant growth regulation in Arabidopsis. Developmental cell, 19(5), 765–777.

Sun, Y., Han, Z., Tang, J., Hu, Z., Chai, C., Zhou, B., & Chai, J. (2013). Structure reveals that BAK1 as a co-receptor recognizes the BRI1-bound brassinolide. Cell research, 23(11), 1326–1329.

Sun, Y., Li, L., Macho, A. P., Han, Z., Hu, Z., Zipfel, C., Zhou, J.-M., & Chai, J. (2013). Structural basis for flg22-induced activation of the Arabidopsis FLS2-BAK1 immune complex. Science, 342(6158), 624–628.

Swings, J., Van den Mooter, M., Vauterin, L., Hoste, B., Gillis, M., Mew, T., & Kersters, K. (1990). Reclassification of the Causal Agents of Bacterial Blight (Xanthomonas campestris pv. oryzae) and Bacterial Leaf Streak (Xanthomonas campestris pv. oryzicola) of Rice as Pathovars of Xanthomonas oryzae (ex Ishiyama 1922) sp. nov., nom. rev. International Journal of Systematic and Evolutionary Microbiology, 40(3), 309–311.

Tina, K., Bhadra, R., & Srinivasan, N. (2007). PIC: protein interactions calculator. Nucleic acids research, 35(suppl_2), W473–W476.

Van Der Spoel, D., Lindahl, E., Hess, B., Groenhof, G., Mark, A. E., & Berendsen, H. J. (2005). GROMACS: fast, flexible, and free. Journal of computational chemistry, 26(16), 1701–1718.

Volkov, A. G., & Ranatunga, D. R. A. (2006). Plants as environmental biosensors. Plant signaling & behavior, 1(3), 105–115.

Wang, G.-L., Song, W.-Y., Ruan, D.-L., Sideris, S., & Ronald, P. C. (1996). The cloned gene, Xa21, confers resistance to multiple Xanthomonas oryzae pv. oryzae isolates in transgenic plants. Molecular plant-microbe interactions: MPMI, 9(9), 850–855.

Wang, H., Chevalier, D., Larue, C., Cho, S. K., & Walker, J. C. (2007). The protein phosphatases and protein kinases of Arabidopsis thaliana. The Arabidopsis Book/American Society of Plant Biologists, 5.

Wang, J., Kucukoglu, M., Zhang, L., Chen, P., Decker, D., Nilsson, O., Jones, B., Sandberg, G., & Zheng, B. (2013). The Arabidopsis LRR-RLK, PXC1, is a regulator of secondary wall formation correlated with the TDIF-PXY/TDR-WOX4 signaling pathway. BMC plant biology, 13(1), 1–11.

Wang, L., Albert, M., Einig, E., Fürst, U., Krust, D., & Felix, G. (2016). The pattern-recognition receptor CORE of Solanaceae detects bacterial cold-shock protein. Nature plants, 2(12), 1–9.

Zhang, X. (1998). Leucine-rich repeat receptor-like kinases in plants. Plant Molecular Biology Reporter, 16(4), 301–311.

Zhang, X. S., Choi, J. H., Heinz, J., & Chetty, C. S. (2006). Domain-specific positive selection contributes to the evolution of Arabidopsis leucine-rich repeat receptor-like kinase (LRR RLK) genes. Journal of molecular evolution, 63(5), 612–621.

