## Supplementary for "Effect of D128N mutation on OsSERK2 in Xa21 mediated immune complex: an *in-silico* study"

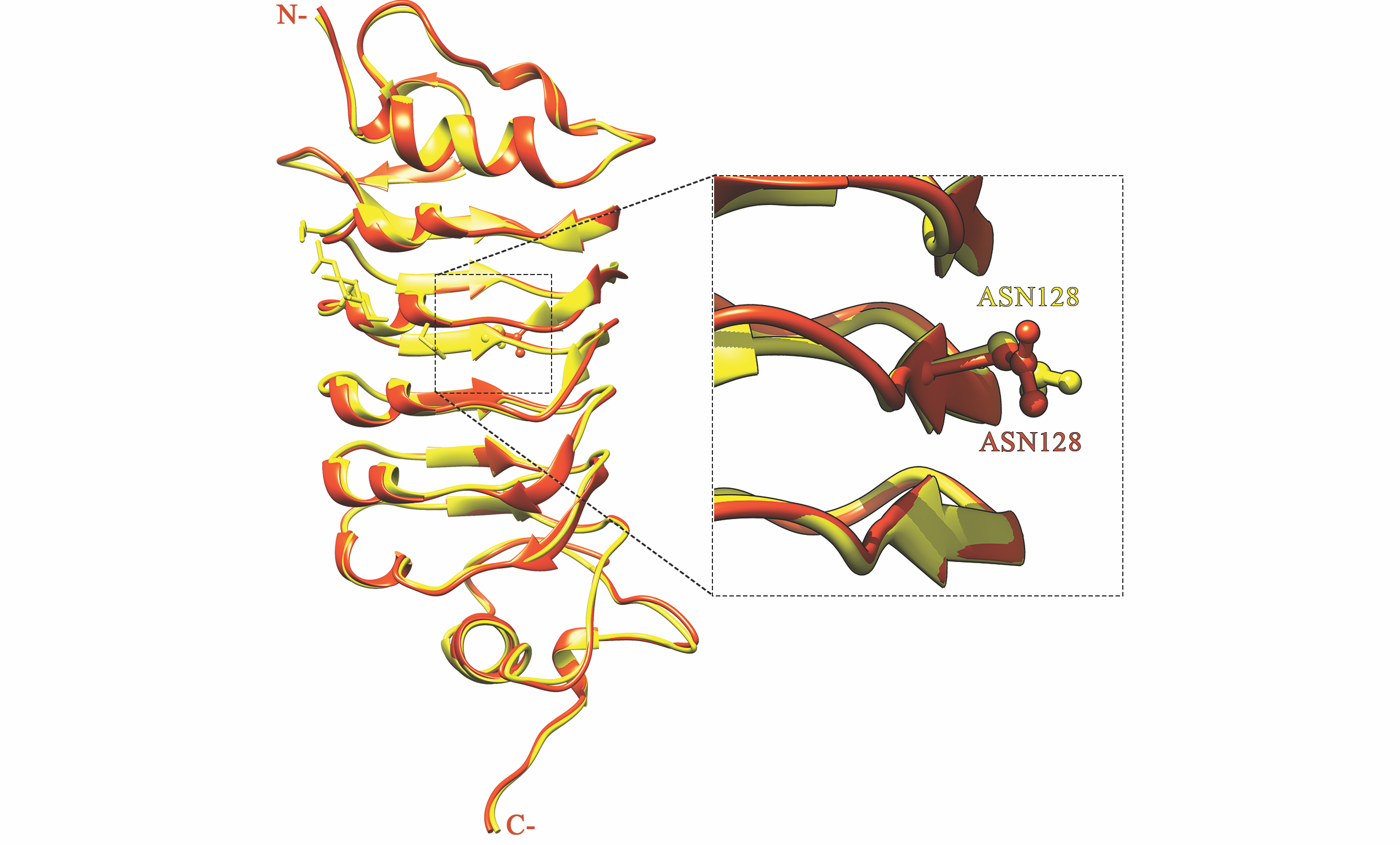

**Fig S1. Superposed structure of our designed OsSERK2 and the crystal OsSERK2 D128N (4q3i).** The crystal OsSERK2 D128N structure is shown in cartoon red structure, whereas the designed structure is shown in cartoon orange. For both cases, the mutated residue is shown in detail with the ball and stick model of Chimera visualization software.

**Table S1.** Salt bridge interaction of different residues from OsSERK2 D128N LRR.

| Atom | Residue | Position | Atom | Residue | Position | Distance |
| --- | --- | --- | --- | --- | --- | --- |
| NH1 | ARG | 78 | OD2 | Asp | 80 | 3.04 |
| NH1 | ARG | 78 | OE1 | Glu | 104 | 3.11 |
| NH1 | ARG | 152 | OE1 | Glu | 174 | 3.75 |
| NH4 | HIS | 202 | OD1 | Asp | 178 | 3.23 |
| NH1 | ARG | 212 | OD2 | Asp | 187 | 3.26 |
| NH2 | ARG | 218 | OD2 | Asp | 214 | 3.35 |

**Table S2.** Side Chain-Side Chain Hydrogen Bonds between Xa21 LRR (chain A), RaxX21-sY (chain B) and OsSERK2 D128N LRR (chain C) within 3 Angstroms

|  | **DONOR** |  |  |  | **ACCEPTOR** | |  |  |
| --- | --- | --- | --- | --- | --- | --- | --- | --- |
| **POS** | **CHAIN** | **RES** | **ATOM** | **POS** | **CHAIN** | **RES** | **ATOM** | **Distance** |
| 231 | A | GLU | OE1 | 53 | C | GLN | NE2 | 2.76 |
| 231 | A | GLU | OE1 | 53 | C | GLN | NE2 | 2.76 |
| 567 | A | THR | OG1 | 140 | C | GLU | OE2 | 2.97 |
| 15 | B | LYS | NZ | 383 | A | ASN | OD1 | 2.99 |
| 53 | C | GLN | NE2 | 231 | A | GLU | OE1 | 2.76 |
| 53 | C | GLN | NE2 | 231 | A | GLU | OE1 | 2.76 |
| 230 | A | ARG | NH2 | 56 | C | ASP | OD2 | 3.31 |

**Table S3.** Hydrophobic Interactions between Xa21 LRR (chain A), RaxX21-sY (chain B) and OsSERK2 D128N LRR (chain C) within 5 Angstroms

| **Position** | **Residue** | **Chain** | **Position** | **Residue** | **Chain** |
| --- | --- | --- | --- | --- | --- |
| 8 | PRO | B | 39 | TYR | C |
| 10 | PRO | B | 37 | ALA | C |
| 88 | TYR | A | 77 | ILE | C |
| 183 | PHE | A | 59 | LEU | C |
| 303 | TRP | A | 19 | PRO | B |
| 354 | PHE | A | 2 | VAL | B |
| 404 | LEU | A | 14 | PRO | B |
| 426 | ILE | A | 14 | PRO | B |
| 428 | LEU | A | 14 | PRO | B |
| 568 | MET | A | 139 | PRO | C |
| 588 | ALA | A | 137 | PHE | C |
| 591 | ALA | A | 137 | PHE | C |

**Table S4.** Measuring the score of the complexes before and after relaxation using Rosetta

| Parameter | Xa21-RaxX21-sY-OsSERK2 | | Xa21-RaxX21-sY-OsSERK2 D128N | |
| --- | --- | --- | --- | --- |
|  | Before Relax | After Relax | Before Relax | After Relax |
| total_score | 8917.146 | 1795.320 | 13230.945 | 1685.627 |
| dslf_fa13 | -4.564 | -3.847 | -4.757 | -5.166 |
| fa_atr | -4616.443 | -4747.295 | -4650.156 | -4728.319 |
| fa_dun | 2528.104 | 885.835 | 2580.683 | 857.742 |
| fa_elec | -1166.172 | -1268.148 | -1140.209 | -1261.704 |
| fa_intra_rep | 10.022 | 9.411 | 10.113 | 10.453 |
| fa_intra_sol_xover4 | 146.113 | 142.103 | 149.389 | 149.451 |
| fa_rep | 7865.523 | 4366.934 | 11732.647 | 4359.160 |
| fa_sol | 2884.582 | 2702.548 | 2930.510 | 2707.819 |
| hbond_bb_sc | -147.812 | -217.407 | -133.045 | -204.902 |
| hbond_lr_bb | -212.549 | -263.964 | -221.640 | -264.163 |
| hbond_sc | -72.987 | -107.682 | -46.856 | -110.765 |
| hbond_sr_bb | -142.591 | -176.260 | -158.294 | -176.639 |
| linear_chainbreak | 0.000 | 0.000 | 0.000 | 0.000 |
| lk_ball_wtd | -51.586 | -103.976 | -66.342 | -103.918 |
| omega | 465.023 | 137.614 | 511.297 | 132.918 |
| overlap_chainbreak | 0.000 | 0.000 | 0.000 | 0.000 |
| p_aa_pp | -75.203 | -166.117 | -73.336 | -174.386 |
| pro_close | 675.120 | 95.876 | 1034.778 | 35.955 |
| rama_prepro | 544.461 | 222.073 | 488.174 | 175.026 |
| ref | 285.995 | 285.995 | 286.800 | 286.800 |
| yhh_planarity | 2.111 | 1.628 | 1.187 | 0.264 |
